# Evaluating genotyping strategies for a small managed population with simulation

**DOI:** 10.1101/2025.01.23.634495

**Authors:** Audrey AA Martin, Jeffrey Schoenebeck, Dylan N. Clements, Tom Lewis, Pamela Wiener, Gregor Gorjanc

## Abstract

**Background:** Collecting genomic information is crucial to advance breeding for complex traits such as health, welfare, and behaviour in domesticated populations. For that purpose, different data collection scenarios can be envisioned based on the number of individuals, the number of markers, and the genotyping technology. This study developed a simulation framework, based on a service dog population, aiming to identify an optimal and cost-effective genotyping strategy that would support the implementation of genomic selection, investigation of the genetic architecture of traits of interest, and track loci of interest.

**Methods:** We simulated a population based on the existing pedigree, using the gene drop method in AlphaSimR. The existing pedigree was extended with additional progeny generations to evaluate the outcomes of different genotyping strategies in the future. We generated genotype data based on existing high-coverage whole-genome sequences (WGS) for the current breeding dogs and evaluated different scenarios for genotyping the progeny. The genotyping options considered SNP arrays of various densities and WGS callsets produced from different sequencing depths. We then phased and imputed the genotype data to high-coverage WGS using AlphaPeel.

**Results:** All scenarios were compared based on individual imputation accuracy against the simulated true whole-genome genotype. Averaged over five generations of simulated progeny, low-pass sequencing (0.5 to 2X depth) achieved accuracies of 0.998 to 0.999. The accuracy of SNP array genotyping (25K to 710K markers) was lower, with means of 0.911 to 0.938.

**Conclusions:** Our simulation was tailored to identify the most cost-effective and efficient strategy for downstream use in genomic selection and genetic research into traits and loci of interest. Low-pass sequencing outperformed SNP array genotyping in imputation accuracy of whole-genome genotypes as expected. Additionally, low-pass sequencing technology was the most affordable genotyping approach currently available for dogs. Thus, it appears to be the optimal choice for balancing the goals of regimented breeding programmes such as those that produce service dogs. This simulation framework could also be adapted to address other objectives for breeding organisations working with small populations.

## Background

Since its introduction, the application of genomic selection has enabled more efficient breeding for heritable traits [1]. Over the last 15 years, the advantages of using genomic selection over traditional selection have been reported for many agricultural plant and animal species (e.g. [2, 3]). However, implementing genomic selection requires a large training population and a coordinated system for evaluations [4]. Populations with a small breeding population size or highly stratified structure could benefit from genomic selection to efficiently reach their breeding goals, yet requirements for its implementation are difficult to satisfy for these types of populations. In these populations, selection is decentralised by ownership, and thus work towards large-scale breeding objectives, such as improving health within a breed, rely on voluntary participation and incentives.

Furthermore, a major challenge to the application of genomic selection in small populations is the trade-off between potential gains and costs. Obtaining a sufficiently large training population to provide accurate genomic predictions requires most, if not all, of the population to have genomic information. Therefore, the required investment to collect sufficient information is expensive, and the initial set-up is labour-intensive. Once established, genomic selection has been proven to be fruitful and to result in substantial returns on investment [1]. However, breeding organisations that manage small populations are generally limited in their resources. Hence, genomic selection represents a major long-term investment for their breeding programs and approaches that reduce the costs of generating genomic information are particularly valuable.

This project created a framework to test genotyping strategies in small managed populations, which can then be adapted to the breeding organisations’ goals, building on previous studies about genotyping strategies (e.g., [5, 6, 7, 8, 9]). We based our study on the case of a service dog population, the UK Guide Dogs (GD) for the Blind Association population, where the long term goal is to improve their population using genomic information. In contrast to other dog populations, service dog populations may be more amenable to genomic selection due to these organisations centralisation and control over breeding. Over decades of phenotypic selection, and more recently pedigree-based selection, these organizations generally have compiled valuable phenotype and pedigree records to improve behavioural suitability and health of their animals. However, many service dog organisations and other breeding organisations managing small populations lack the genomic information that would allow implementation of genomic selection.

There are several methods for collecting genomic information, each with its own advantages and limitations for a population; based on the number of individuals, markers, genotyping technology, traits and loci of interest, and budget. The three main approaches are SNP genotyping, high-coverage whole-genome sequencing (WGS), and low-coverage WGS (also known as skim or low-pass sequencing). SNP genotyping, the most established method in selective breeding, is cost-effective and provides manageable data. This approach captures genome-wide information, supporting genomic selection, but lacks resolution for fine-mapping within genome-wide association studies. Additionally, SNP arrays are designed based on prior knowledge based on specific populations, limiting the capture of novel variation specific to other populations. High-coverage WGS offers high resolution and comprehensive genetic analysis, with the capacity to capture all types of variation (coverage- and technology-dependent), yet it is time-, computationally-, and storage-intensive, and is generally costly. Although it accurately captures a smaller portion of the entire genome, low-pass sequencing may provide a more affordable option, making it more suitable for some circumstances. In combination with targeted high-coverage WGS and accurate genotypic imputation, the low-pass technology may provide sufficient resolution across the genome.

Simulation allows a comparison of these approaches before application to the population, supporting strategic decision-making. To evaluate the different genotyping strategies in this project, we developed a method using the AlphaSimR [10] to simulate a population based on the GD population, using their pedigree and existing high-coverage WGS data. The genotyping strategies were compared based on the following criteria: 1) data for potential downstream use in genomic selection, 2) costs, 3) data processing load, and 4) utility of data to improve the understanding of the dog genome and the genetic architecture of traits and loci of interest.

## Material and Methods

### Pedigree information

The GD pedigree on which simulations were based consisted of 62,698 individuals, 32,607 females and 30,091 males, spread across 31 generations from 1901 to 2023. More than 80% of the population was born after 1975. There were 5,683 full founders, individual with both parents unknown, and 1,088 half founders, individual with only one parent known, spread throughout the generations of the pedigree, giving 6,771 founders in total. On average, individuals had complete information for 4.4 generations (ranging from 0 to 9) and partial information for 16.6 generations (ranging from 0 to 30). The majority of the population was composed of three main breeds or their crossbreeds: Labrador Retriever, with 23,901 purebreds and 20,369 crosses; Golden Retriever, with 8,612 purebreds and 21,854 crosses; and German Shepherd Dogs, with 5,035 purebreds and 947 crosses (see Additional file 1). In total, there were 22 known breeds present in the pedigree. Additionally, coverage information on recently generated WGS data for the current breeding dogs (215 females and 70 males) was also used in the simulation (see Additional file 2).

### Simulation

The aim was to simulate a breeding population based on that of a service dog population. An R script using AlphaSimR [10] was created to produce the overall simulation pipeline. Information on numbers of founders and the current breeding population matched the GD pedigree. The following steps were then implemented for the simulation:

- Set-up of genome parameters,
- Creation of SNP arrays,
- Generation of the current population,
- Creation of future generations,
- Creation of SNP marker genotype and WGS datasets, and
- Imputation.

The set-up of the genome parameters and the creation of the SNP arrays were performed once and used across replicates of gene dropping within pedigree to focus on imputation accuracy within the pedigree.

#### 1. Set-up of genome parameters

Based on the number of SNP and indels found in a preliminary analysis of WGS for 285 GD breeding dogs, we assumed a total of 250,000 segregating sites for *Canis lupus familiaris* autosome (CFA) 1. Other simulation parameters were based on CFA1 information reported by the PopSim consortium [11, 12]: physical length of 122Mb, genetic length of 90cM (based on recombination rate of 0.74*×*10*^−^*^8^). However, due to computational constraints of running several replicates and scenarios, we simulated only a fragment of CFA1 containing 10,000 biallelic segregating sites (10,000/250,000 = 0.04 of CFA1), corresponding to 4.91Mb and 3.7cM.

#### 2. Creation of SNP arrays

As for the chromosomal parameters mentioned above, the number of SNPs simulated on the chromosome was scaled by 0.04. We simulated the following SNP genotype arrays as follows:

- 710,000 SNP array (Affymetrix-like):
  **–** 18,205 SNP estimated for a simulated chromosome (710,000/39 canine chromosomes).
  **–** 728 SNP simulated (=18,205 * 0.04)

- 170,000 SNP array (Illumina-like):
  **–** 4,360 SNP estimated for a simulated chromosome.
  **–** 174 SNP simulated.

- 50,000 SNP array:
  **–** 1,280 SNP estimated for a simulated chromosome.
  **–** 51 SNP simulated.

- 25,000 SNP array:
  **–** 640 SNP estimated for a simulated chromosome.
  **–** 26 SNP simulated.

The two first arrays were designed following real arrays: the Affymetrix-like mimics the Axiom^TM^ Canine HD Array [13] and the Illumina-like mimics the CanineHD BeadChip [14]. To our knowledge, there is currently no lower density array commercially available for dog genotyping. Nonetheless, we assessed the theoretical capabilities of 50K and 25K arrays, as might be produced by custom designs. The highest density array was created first. We generated a dummy GD population using *quickHaplo* and *pedigreeCross* functions in AlphaSimR, as described in more detail in the following sections. We then extracted all segregating sites from the genomes, and randomly selected 728 sites uniformly across the allele frequency distribution. Markers for lower density arrays were then randomly sampled from previously created arrays, e.g. the 170K array was sampled from the 710K array. Thus the smaller arrays were nested within the larger ones. The arrays were created once and commonly shared between all five replicates.

#### 3. Generation of the current population

Using the *quickHaplo* function within AlphaSimR, we created a population of 6,771 founders (including both full and half founders), each with a genome fragement with 10,000 loci for which alleles were randomly assigned. We then performed a gene drop simulation using an adapted version of *pedigreeCross* function in AlphaSimR. The haplotypes were dropped through the generations with recombination, by mating all individuals as per the pedigree information, starting from the founders. Half-founders inherited one genome from the known parent and the other from the founder population to represent the unknown parent. In this step, true genotypes and haplotypes for the whole simulated population were created and stored.

#### 4. Creation of future generations

For the creation of future generations, we simulated a breeding scheme, represented in Figure 1, mimicking the population of interest that will transition to genomic selection. For each of the following five years, we simulated a new generation of progeny. In the first two years, individuals from the current breeding population were mated to produce a litter of 8 puppies, as per the average within our case population. All 215 females were mated, and the 70 males were mated between 0 to 4 times with probability 1%, 4%, 10%, 55%, and 30%, respectively. From year 3 onwards, the size of the breeding population was increased to 240 for the females and 80 for the males, to reach 1,920 progeny per year, mimicking a small population under genomic selection (with a higher number of young individuals and a higher replacement rate). From the current breeding stock (Gen 0), 35 of the 70 males and 60 of the 215 females were retired and replaced by selected two-year old (born two generations before) individuals, old enough to be consider as a breeding candidate. Thereafter, every year, part of the breeding population was retired (85 out of 240 for the females and 45 out of 80 for the males) and replaced by new breeding dogs, selected from the list of candidate progeny. Over time, the current breeding population with high-coverage WGS data was entirely replaced by breeding dogs with genomic information, as specified in the genotyping scenarios described below. Decisions for the mating program and the composition of the breeding population were simulated as random due to the absence of phenotype data in our case study. Although not realistic, we do not expect our results to be affected, as we are comparing the different scenarios based on imputation accuracy (described below) rather than genomic prediction accuracy.

**Figure 1.**
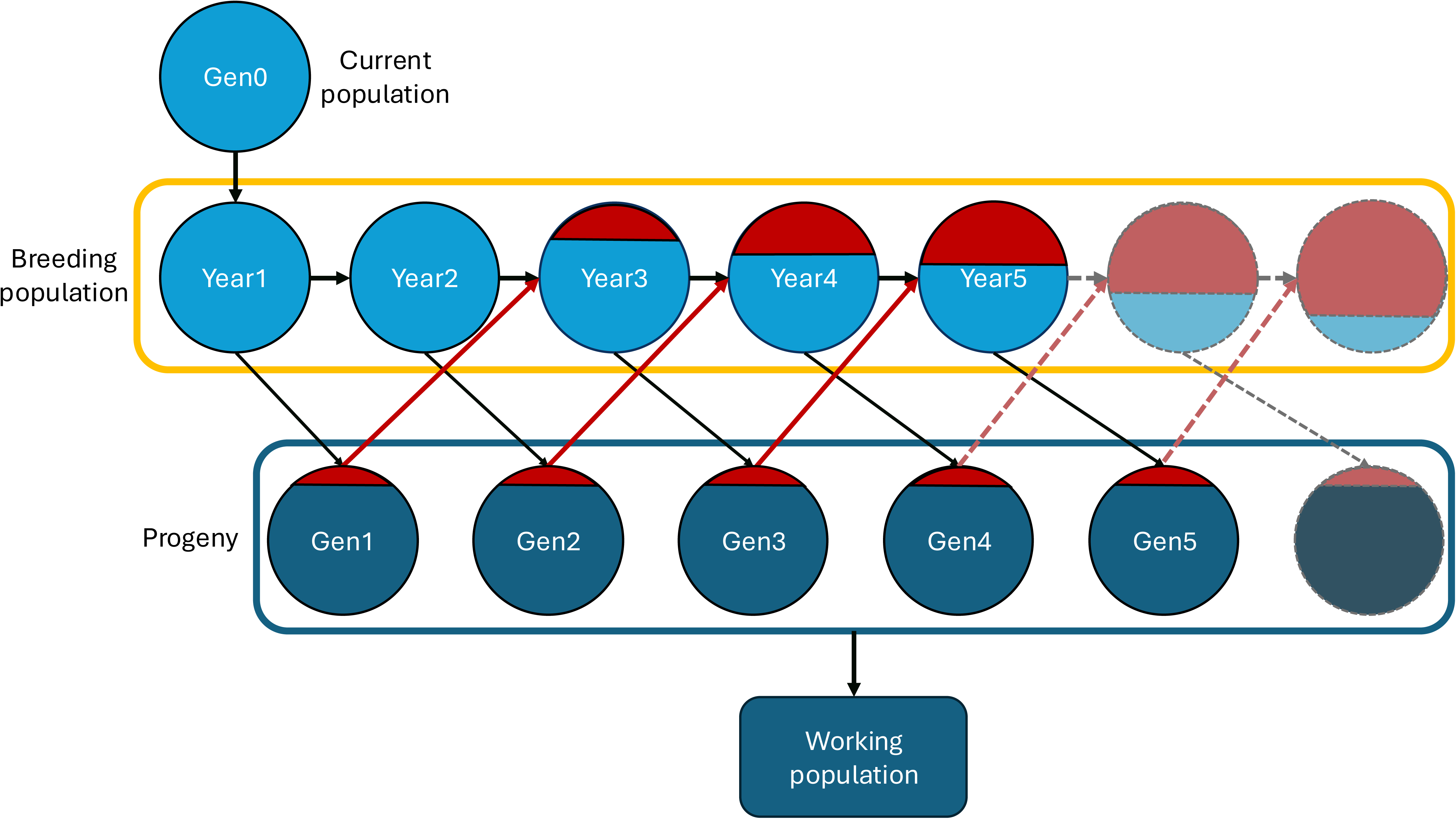
Simulated breeding scheme for a numerically small population transitioning from pedigree-based selection to genomic selection. Timeline of the simulated breeding program, spanning over five new generations (Gen1 to Gen5), with potential continuation beyond Year 5 illustrated by dashed and muted-coloured elements. During the first two years, the breeding population is exclusively composed of the current breeding individuals (Gen0). From Year 3 onwards, a subset of progeny from two years prior (at sexual maturity) is selected to renew the breeding population (red elements). Arrows illustrate the flow of individuals between groups, representing the structured progression of the breeding program.

#### 5. Creation of SNP marker genotype and WGS datasets

*Approaches* Multiple scenarios were simulated to test diverse genotyping approaches. First, from the true genotypes and haplotypes described above, WGS data for the current breeding dogs was generated, with coverage matching the empirically observed WGS coverages from GD breeding dogs (Additional file 2). Secondly, genotype data for the progeny was simulated based on various approaches: 20X coverage sequencing, low-pass sequencing (0.5 to 2X), and SNP array genotyping. A negative control, where no genomic information was collected for the progeny, was also performed.

For the low-pass sequencing scenarios, we added an additional option (“amp”, for amplification), where small targeted genomic regions (loci) were sequenced at a 30X depth. The objective was to determine if using amplification for locus of interest, such as known causal mutations for disease, increased the individual marker accuracy of that locus compared to low-pass scenarios without amplification. For the scenarios with this option, we simulated one amplified locus of interest, placed at random on the chromosome and common to all replicates.

For the low-pass sequencing and SNP array genotyping scenarios, we also included a further approach “addseq”, where progeny selected as breeding dogs for subsequent generations were sequenced at 20X coverage. The aim of this approach was to improve the quality of the genomic information available overall, as the breeding population would be continuously composed of dogs with high-coverage WGS data. The 22 scenarios listed in Table 1, incorporated for each of the 5 replicates, employed various approaches to collect genomic information on the progeny.

**Table 1.** Summary of the 22 genotyping scenarios tested with simulation for a small population. There are two baseline scenarios:“Gold standard” where high-coverage WGS for the whole population is available, and “None” where only genomic information for current breeding individuals was used, without additional genomic information collected on the progeny.“amp” represents the use of the amplification technology (30X) for the locus of interest. The parent population always contained the current breeding individuals with high-coverage WGS data matching real data. For the “addseq” scenarios, progeny selected as breeding individuals were additionally sequenced at 20X, ensuring high-coverage sequencing for the entire breeding population.

|  | Scenarios | Parents Sequencing |  | Progeny Sequencing |  |  |  |  | SNP array genotyping |  |  |  |
| --- | --- | --- | --- | --- | --- | --- | --- | --- | --- | --- | --- | --- |
|  |  | Current individuals | Addseq | 20X | 2X | 1X | 0.5X | amp | 710K | 170K | 50K | 25K |
| Baseline Scenarios | None | X |  |  |  |  |  |  |  |  |  |  |
|  | Gold standard | X |  | X |  |  |  |  |  |  |  |  |
| Low-pass sequencing scenarios | 2X | X |  |  | X |  |  |  |  |  |  |  |
|  | 2Xamp | X |  |  | X |  |  | X |  |  |  |  |
|  | addseq_2X |  | X |  | X |  |  |  |  |  |  |  |
|  | 1X | X |  |  |  | X |  |  |  |  |  |  |
|  | 1Xamp | X |  |  |  | X |  | X |  |  |  |  |
|  | addseq_1X |  | X |  |  | X |  |  |  |  |  |  |
|  | 0.5X | X |  |  |  |  | X |  |  |  |  |  |
|  | 0.5Xamp | X |  |  |  |  | X | X |  |  |  |  |
|  | addseq_0.5X |  | X |  |  |  | X |  |  |  |  |  |
| SNP array genotyping scenarios | 710K | X |  |  |  |  |  |  | X |  |  |  |
|  | addseq_710K |  | X |  |  |  |  |  | X |  |  |  |
|  | 170K | X |  |  |  |  |  |  |  | X |  |  |
|  | addseq_170K |  | X |  |  |  |  |  |  | X |  |  |
|  | 50K | X |  |  |  |  |  |  |  |  | X |  |
|  | addseq_50K |  | X |  |  |  |  |  |  |  | X |  |
|  | 25K | X |  |  |  |  |  |  |  |  |  | X |
|  | addseq_25K |  | X |  |  |  |  |  |  |  |  | X |

*Generation of SNP array genotype files* The creation of the SNP array genotype files were based on the simulated true genotypes. Coded as 0 (homozygous for reference allele), 1 (heterozygous), 2 (homozygous for alternate allele), and 9 (missing), genotypes were extracted from the WGS data at marker positions for the described SNP arrays. Errors were then added to mimic SNP array genotyping. First, using a binomial distribution, the probability of observing an error at a locus was set to 1*e^−^*^4^ [15, 16]. There were then three different types of error possible:

- a single-allele change, (genotype 0 to 1, 1 to 0, 1 to 2, or 2 to 1), with a probability of 1 *−* 1*e^−^*^5^ *−* 5*e^−^*^5^ = 0.99994,
- a double-allele change, (genotype 0 to 2 or 2 to 0), with a probability of 1*e^−^*^5^,
- missing information with a probability of 5*e^−^*^5^.

*Generation of WGS files* WGS data was also extracted from the simulated true whole-genome sequence information. There were four steps to create the WGS data:

- *1) Locus-specific sequenceability* We incorporated the fact that some loci either generate more or less sequence reads or that the resulting reads are more or less often aligned to the reference genome. Each locus-specific sequenceability represented both the sequencing variability, due its base-specific composition and the read mappability. For the whole genome at the population level, each locus received a “sequenceabality score” defined by a gamma distribution of shape 4 and rate 0.25, as described in [17, 18].
- *2) Number of locus reads per individual* To simulate the random amplification of DNA fragments associated with the sequencing process, we used a Poisson distribution with mean equal to the chosen sequencing depth (20X, 2X, 1X, or 0.5X) multiplied by the locus-specific sequenceability score to calculate the locus-specific sequencing depth. Thus, different numbers of sequence reads per locus per individual were obtained. For the amplified locus (“amp”), we added 30 reads to the corresponding site to achieve 30X amplification.
- *3) Allele reads per individual* For each sequencing read, we attributed either the reference or alternative allele based on the true genotype of the individual at that locus. We used the genotype dosage, count of reference alleles at the locus, to determine the probabilities of the reference (*p_li_*) and alternative ((1 *− p_li_*)) alleles at locus *l* for individual *i*. We sampled the allele read from a binomial distribution with probability (1 *− p_li_*) for the alternative allele.
- *4) Error* To simulate some sequencing errors, the allele present on the read was flipped to the opposite allele with probability 1*e^−^*^3^.

#### 6. Imputation

Once the population was simulated and the data files created, we performed imputation to WGS using AlphaPeel, via multi-locus peeling, a pedigree-based method that iteratively loops though the generations to phase and impute the allele information across the pedigree [19]. This method also uses pedigree and linkage information, by assuming a high probability of alleles at nearby loci to be inherited jointly from the same parental haplotype.

The quality of the imputation was evaluated by estimating correlations between the true simulated genotypes and the imputed genotype dosages across the whole-genome (all segregating sites). We calculated the correlation per individual across markers and then averaged it across individuals. We also determined the average imputation accuracy at the locus of interest across individuals by estimating a marker imputation accuracy, i.e. average of marker accuracy across all individuals, for a given locus. Unless specifically stated otherwise, the accuracy stated refers to individual imputation accuracy. Due to computational constraints, a single final imputation was performed with the information from all generations, rather than imputation within each generation, although in practice, the latter would be more realistic for an actual data collection procedure.

To ensure that the simulated chromosome subset was representative, we compared the imputation results of two scenarios (2X low pass WGS and 710K SNP array) for two independent fragments of 10,000 loci. We expect similar accuracies for the two simulation runs, indicating that the smaller fragment is representative of an entire chromosome, based on the logic that performing imputation on a genomic segment instead of a complete chromosome only reduces the computational load and does not affect the accuracy results. The average imputation accuracies from these tests were not statistically different (Welch two-sample t-test, p-value *>*0.6). Therefore, we concluded that simulating one fragment of 10,000 loci was representative of the whole chromosome.

## Results

For all scenarios, we used the multi-locus peeling method for the imputation to leverage the pedigree and linkage information to propagate the high-density genomic information throughout the population. We first investigated two baseline scenarios. Under the “gold standard” scenario, where the whole population was sequenced at high coverage, we obtained an imputation accuracy close to one, as shown in Additional file 3. In the “None” scenario where only the current breeding dogs (Gen 0) had genomic data, the imputation accuracy for the progeny dropped drastically, as increasing generations separated the new individuals from Gen 0 (detailed in Additional file 3).

Within the SNP array genotyping scenarios, the results depended on the density of the simulated SNP array (Figure 2). The overall imputation accuracy across the five generations of progeny decreased with array density: 0.938 (710K), 0.920 (170K), 0.911 (50K), and 0.911 (25K). These imputation accuracies started high and slowly declined over the generations; for the higher-density arrays, the trend was nearly stable with a slight decline over generations. As expected, SNP array genotyping did not offer further information for the locus of interest since we did not explicitly account for the causative variant or nearby markers in the array design (Additional file 4).

**Figure 2.**
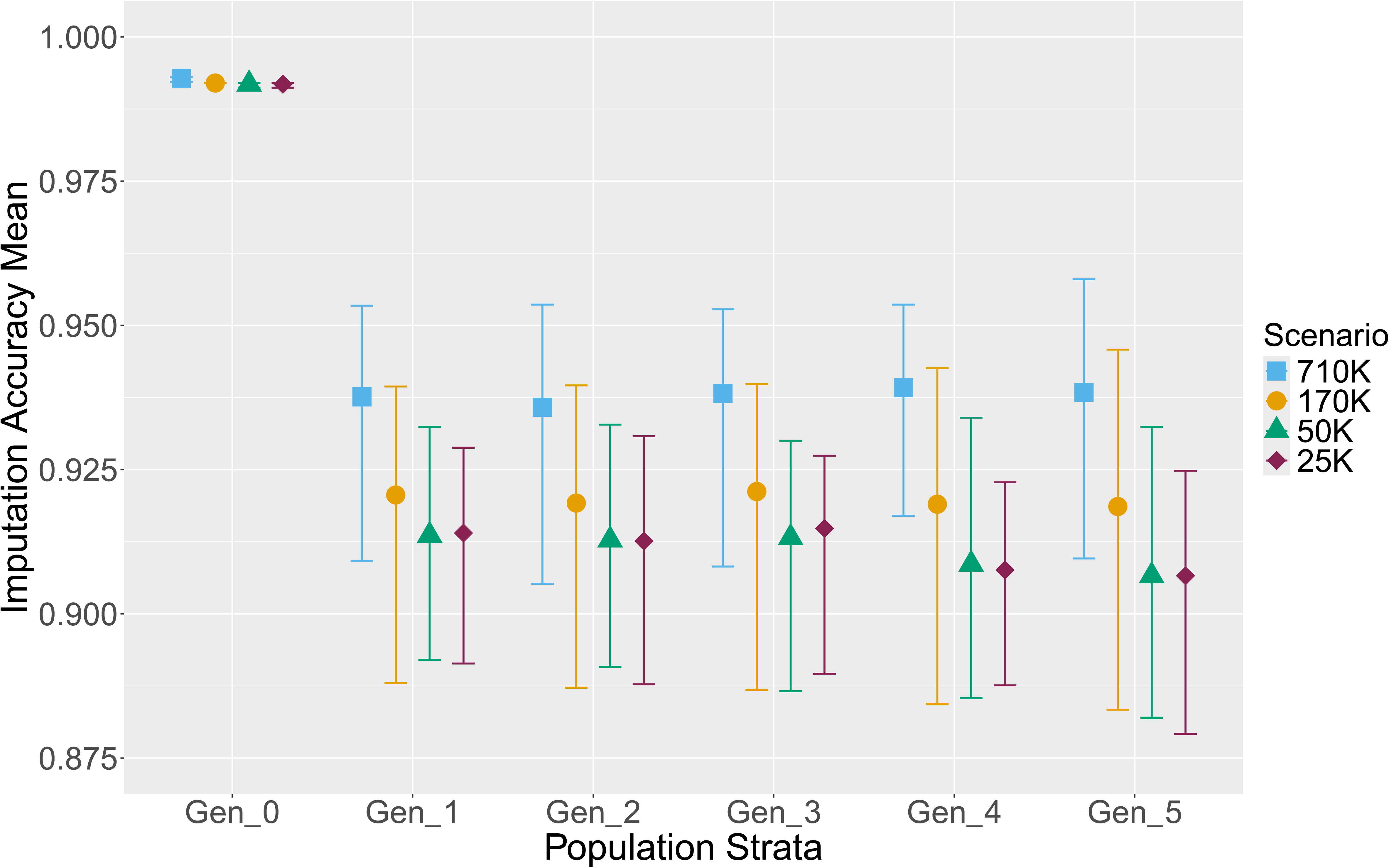
Individual imputation accuracy for SNP array genotyping strategies. The brackets represent the 95% interval for the variation observed across the five simulated replicates.

The other approach using SNP genotyping involved additional 20X coverage WGS data for the breeding population in each generation (“addseq”). In these scenarios, in addition to SNP genotyping, we sequenced genome at 20X coverage for all progeny selected as breeding dogs at two years-old (see red elements in Figure 1). As shown in Figure 3, the addition of sequencing for the new breeding stock was valuable, with average [min : max] gain in imputation accuracy of 0.039 [0.027 : 0.052] for the 710K SNP array, 0.052 [0.035 : 0.070] for the 170K SNP array, 0.059 [0.040 : 0.080] for the 50K SNP array, and 0.058 [0.035 : 0.078] for the 25K SNP array. The difference between the scenarios with and without additional sequencing increased over the generations. By Gen 5, the “addseq” scenarios of high-density SNP genotyping reached an imputation accuracy of at least 0.980.

**Figure 3.**
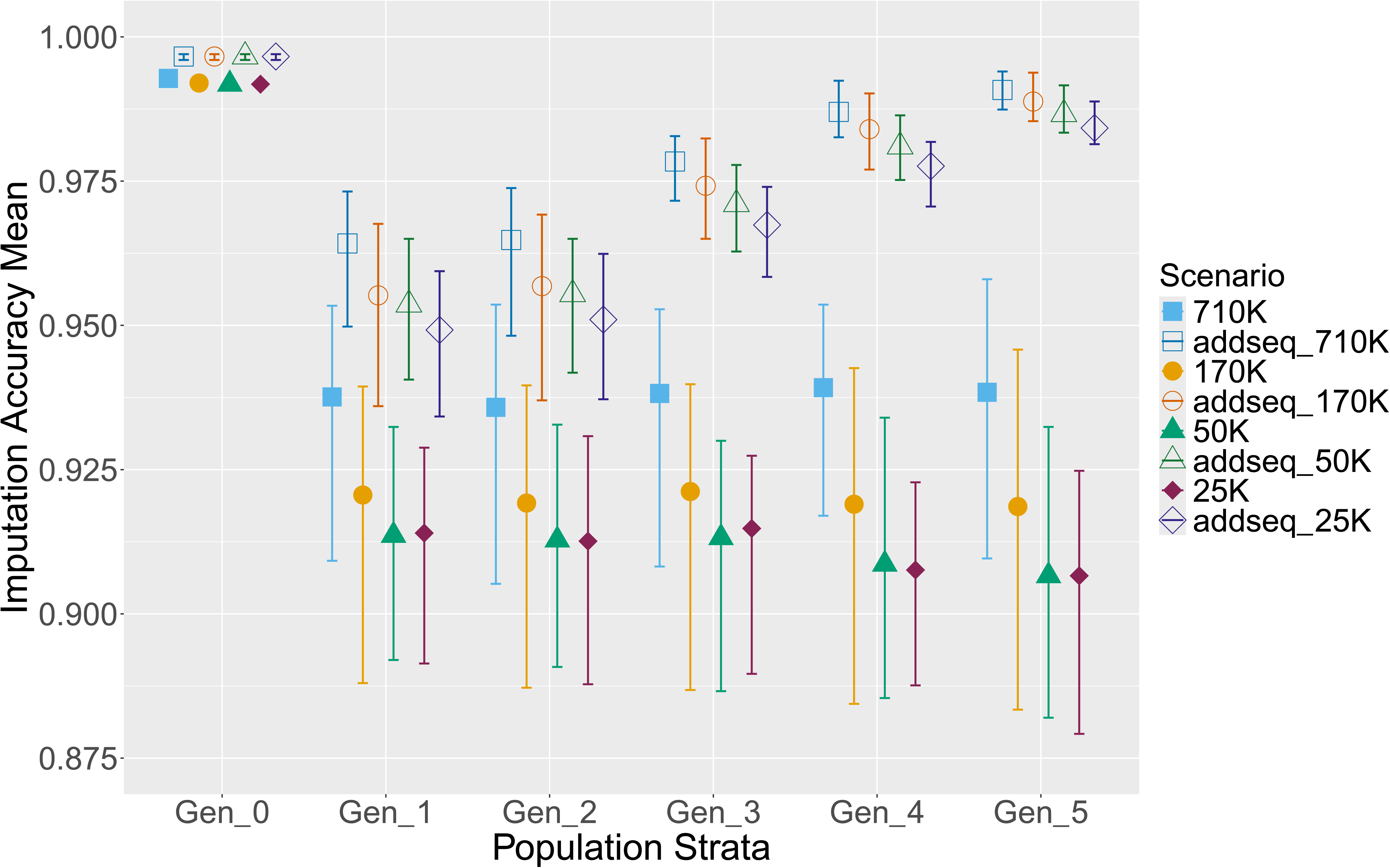
Individual imputation accuracy for SNP array genotyping with additional resequencing (“addseq”) strategies. The brackets represent the 95% interval for the variation observed across the five simulated replicates.

The imputation accuracy for the various low-pass sequencing strategies was much higher than for SNP genotyping, with all averages over 0.995 (Figure 4). Results improved with greater genome sequencing coverage (2X *>*1X *>*0.5X), but all reached accuracies close to 1 with increasing generations. The use of the amplification technology (“amp” scenarios) was partially useful. At the targeted locus of interest, it resulted in marker imputation accuracy close to 1 from the early generations (Additional file 5). However, the amplification did not significantly increase the overall individual imputation accuracy (Figure 4). Similarly, when considering 20X coverage sequencing of the progeny selected for breeding (“addseq” scenarios), we did not observe a strong impact on the imputation accuracy (Additional file 6).

**Figure 4.**
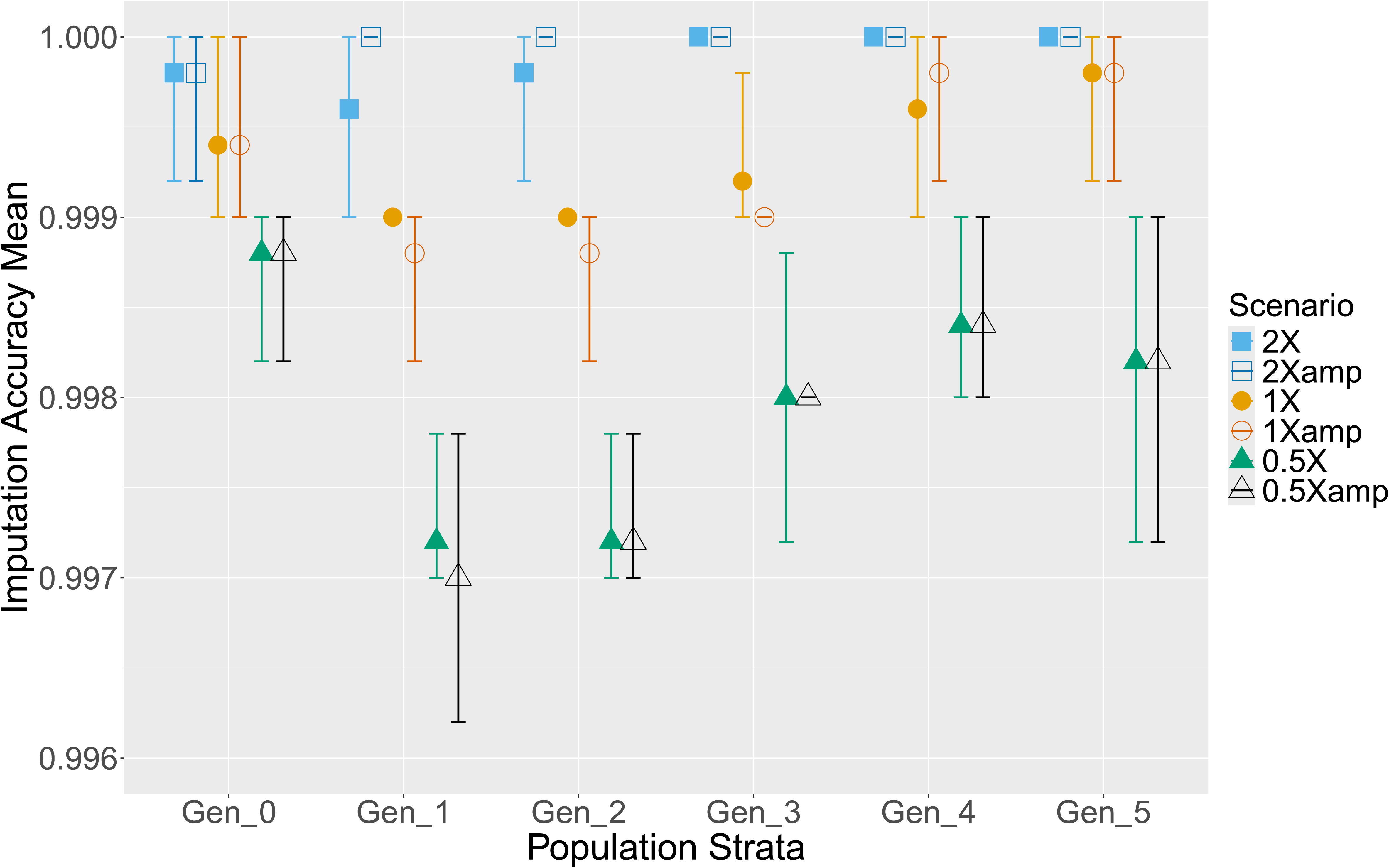
Individual imputation accuracy for low-pass sequencing strategies. The brackets represent the 95% interval for the variation observed across the five simulated replicates.

To more clearly compare scenarios with the most promising results and the most realistic cost-wise for a small breeding organisation (2X, 1X, 0.5X, addseq 710K, and addseq 170K), we plotted them together (Figure 5). Low-pass sequencing, at any coverage, offered maximum imputation accuracy to the WGS level across all generations. Similar results were obtained with SNP genotyping combined with a continuation of 20X coverage sequencing for the breeding population (“addseq”), but high accuracies were only reached in the later generations of the simulation.

**Figure 5.**
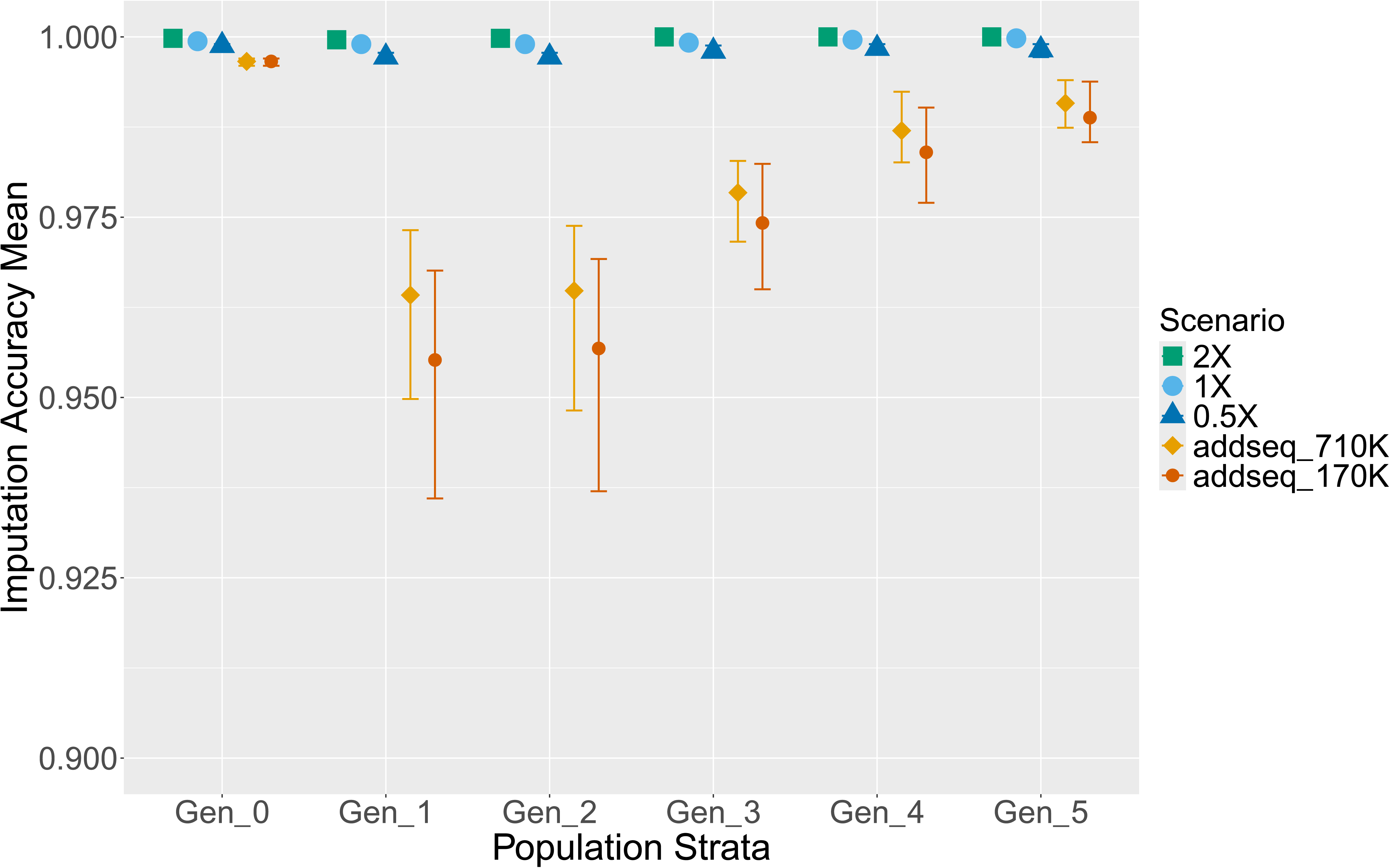
Comparison of the individual imputation accuracy for the best strategies. The brackets represent the 95% interval for the variation observed across the five simulated replicates.

## Discussion

This project created a simulation framework to test strategies to collect WGS information across generations of a breeding population. We based the simulations on the breeding population of Guide Dogs (GD), as the organisation aims to make improvements in the health and welfare of its dogs by use of genomics. The simulation framework was developed to support decision-making by GD, but also has the potential to be applied to other populations. The decision for choosing a strategy for collecting genomic information depends on objectives. In this project, we considered multiple criteria for evaluating the possible genotyping strategies, based on their capacity to: 1) support the implementation of genomic selection, 2) be cost-effective, 3) limit data processing and storage, and 4) provide data to support research and improvement of traits and loci of interest. Below we discuss our results, each of the criteria in turn, and finally the potential of the framework to be applied more widely.

### Imputation accuracy results

The three different depths of low-pass sequencing (2X, 1X, and 0.5X) all yielded high imputation accuracy (*>*0.995) overall and for the locus of interest. There was no benefit to additional sequencing at higher coverage of the progeny selected as breeding dogs (“addseq” scenarios). For the SNP array genotyping, the overall imputation accuracies ranged between 0.900 and 0.950, where accuracy expectedly increased with array density. Over the generations, we noted a slow decrease in imputation accuracy. The marker imputation accuracies for the locus of interest were slightly lower than the individual imputation accuracies, but followed the same slow decline across the generations. Additional sequencing of the selected progeny (“addseq” scenarios) improved the imputation accuracy from the early generations and reversed the decrease such that it exceeded 0.980 for all array densities by Gen 5.

An advantage of sequencing, even at low coverage, over SNP array genotyping, was the collection of information about recombination and mutation at all sites in every generation. AlphaPeel uses pedigree and linkage information to phase and impute genomic data, which benefits from the large litter size and extended generational pedigree information. Over time, the WGS information accumulated and provided an extensive haplotype reference library for imputation across the pedigree. With only SNP array genotyping, the associated haplotype library is also built, but is limited by the SNP array density capturing recombination positions less accurately than sequencing data, which explains the slow decline in imputation accuracy observed over the generations. The addition of 20X coverage sequencing of selected progeny (“addseq” scenarios) to SNP array genotyping resolved this issue by continuing to collect information on the key genomes of the population. In contrast, low-coverage sequencing scenarios did not need this additional sequencing at higher coverage to achieve high imputation accuracy for the whole-genome genotype, as that information was already captured.

It is important to consider that the imputation accuracy values were obtained by simulations based on a simplified version of a service dog population, such that the measured accuracies were likely an overestimation of what would be achieved in practice. Furthermore, for computational reasons, the imputation step was performed once with all five generations in the simulations. However, in actuality, imputation would be performed more regularly, once per generation, which would result in lower imputation accuracy until reaching the same level of information as for a single imputation procedure by the fifth generation. However, these factors should influence all scenarios equally and thus it is valid to compare accuracies between the different scenarios.

### Implications for criteria

#### Genomic selection

A major obstacle to the implementation of genomic selection based on WGS comes from the requirement of an excessively large training population for the accurate estimation of effects for each variant, leading to an unaffordable cost for most breeding organisations. In this study, we considered the case of GD for which only a limited amount of genomic data is available: high-coverage WGS data was produced on their current breeding population. In this case, further data generation is required to assemble a reference population of adequate size. Consequently, we evaluated scenarios collecting information on the whole GD population, with the objective to collect enough genotypes to create an adequate training population for future genomic prediction.

When routine WGS first became a possibility, simulations suggested that the larger and denser set of SNP provided by WGS would increase prediction accuracy [20, 21]. However, this early promise has not been realised; the gains with WGS are reported to be small at best and results are highly trait- and population-dependent as reviewed in [22]. Frequently, populations under intensive selection have a low genetic diversity (as measured by a small effective population size), which can already be captured adequately by SNP array genotyping, leading to limited advantage of WGS [23]. Thus for a simple population, such as a single breed, low to medium density marker arrays are generally sufficient for accurate genomic prediction and subsequent selection in practice [1, 24]. However, for populations containing multiple breeds and different levels of admixture, as in the case of the GD population, higher density or markers selected from WGS can capture more information, including causal variants, and potentially offer higher prediction accuracy [25, 26]. Independent of the genotyping technology and imputation, we could expect a good performance with genomic selection considering Labrador Retrievers, a popular breed for service dogs. For example, based on the method described in [27], averaged estimate of behaviour traits’ pedigree-based heritability [28] (*h*^2^ = 0.19) and effective population size [29] (*Ne* = 86), we predict an accuracy of 0.60 for the genomic estimated breeding values in the Labrador Retriever sub-population of GD (around 1,600 individuals for two generations). Accordingly, all the tested scenarios are likely to allow the implementation of genomic selection with acceptable levels of prediction accuracy. This could be achieved by using either a sufficiently dense SNP array or WGS data, though the latter might require linkage-disequilibrium-based pruning of markers to improve the statistical signal and reduce noise in the genomic prediction model with such population size. However, when considering the whole GD population, prediction accuracy for the many cross-bred individuals may be considerably lower, with the difficulty to estimate accurate genomic values in crossbred and admixed populations (see review in [30]). The use of WGS data, obtained through imputation, may alleviate this issue, but it is important to secure accurate genomic data as genomic prediction accuracy has been shown to decrease as imputation errors accumulate [6, 18, 31]. Therefore, due to the higher imputation accuracy we observed, low-pass sequencing would thus be the preferred option over SNP array genotyping, even with supplementation of WGS in the breeding stock, when using WGS data for future genomic predictions. Additionally, the combination of low-pass sequencing and imputation would offer the advantage of incorporating the functional information as provided by WGS, while keeping the costs down (see below), making the creation of a large training population with accurate WGS data more achievable.

#### Costs

We contacted genotyping providers to obtain cost estimates for genotyping 2000 dogs, for the different scenarios. These estimates should be considered at the level of scale comparison rather than actual values as prices are continuously changing and depend on the genotyping volume. Briefly, the cost of the data generation would be lowest for strategies relying on low-pass sequencing, followed by SNP genotyping. Finally, all scenarios involving 20X coverage sequencing (“gold standard” and “addseq”) would be the most expensive.

The “gold standard” scenario included 20X sequencing for all progeny, for which the cost per dog was quoted as around £300. Although the price of sequencing has been falling over the years, 20X coverage sequencing is still too expensive to be a routine process for most breeding programs. However, at lower coverage, the technology is more affordable and now accessible for large-scale deployment. We estimate the cost of 2X, 1X, and 0.5X sequencing at around £45, £30, and £25 per dog, respectively. Assuming a cost of 10%, 20%, and 30% of the original 1X sequencing price, the amplification technology (“amp” scenarios), sometimes called target capture, would cost an additional £3, £6, and £9, respectively. Note that services providing low-pass sequencing at 1X and imputation to WGS for around £30 per dog are currently available.

SNP array genotyping is a well-established genotyping method routinely used across many livestock species and its populations. In dogs, there are different approaches to the use of this technology. Several providers offer genotyping services to the general public for breed identification and genetic health testing. These genotype analyses are performed based on private arrays and databases and the resulting interpretations for an individual dog are provided to the client. Unlike the case for livestock, genome-wide SNP genotyping in dogs is largely done by research groups, explaining the absence of low-density arrays on the market, and the high costs result in limited scale of genotyping. Both the Affymetrix (710K) and Illumina (170K) arrays were originally developed for genome-wide association studies, covering veterinary research and studies using dogs as a model organism for human disease, and have currently many other applications. These providers offer dense genomic coverage of markers segregating across an extensive set of breeds. Costs for genotyping using the 710K and the 170K arrays have been quoted at around £45 and £70 per dog, respectively. The volume of arrays produced and processed each year by the genotyping provider strongly influences the SNP genotyping cost, explaining the high price estimates for our case population with the low volume of arrays processed in dogs in comparison to livestock species. In genetically small livestock populations, such as local cattle or sheep breeds, such costs can drop under £10 per individual by using popular arrays also used by larger breeds.

Costs for 25K and 50K arrays, which are not currently available commercially for dogs, are more difficult to estimate as they would depend on the cost of design process and genotyping volume. It is feasible to design a customised array aimed at research and applied purposes, although a difficulty may arise from the commercial patenting of causal SNPs for the known monogenic diseases in dogs (see below). Service dog organizations could consider collaborating to design a custom low- or moderate-density SNP array in order to share the costs more widely.

### Data processing

Other major factors for decision-making are the cost and logistics of dealing with genomic data. SNP array genotyping is more practical than sequencing for small organisations as it does not entail extensive memory, data storage, and bioinformatics requirements. However, scenarios with SNP array genotyping performed better in terms of imputation accuracy when combined with 20X coverage WGS data (as measured by the WGS imputation accuracy). Therefore, for such scenarios, both SNP array and WGS data would need to be handled and prepared, when WGS level information is required to incorporate population-specific loci and loci of interest. The pipeline for managing sequence data requires read alignment, multiple quality controls, variant calling, and variant annotation. In addition, the SNP array genotyping data would need to be imputed to WGS level. Thus, for all of the scenarios giving highly accurate data (20X coverage WGS, low-pass sequencing, or SNP array genotyping with “addseq”), computationally-demanding processing pipelines would be involved, which are both time- and storage-demanding and require specific expertise. This presents substantial initial investment and running costs for small breeding organisations. As mentioned above, some providers now offer both low-pass sequencing and subsequent imputation within one service package, which is an opportunity for small organisations to reduce the processing and imputation burdens.

Other providers offer a more bespoke platform for bioinformatics and imputation analyses as well as data storage. Such arrangements would need to be investigated on a case-by-case basis to determine whether this option would be cost-effective for an individual breeding organisation.

### Genetic basis of traits of interest

The history of dog breeds, involving small founding populations, bottlenecks, use of popular sires, and strong selection for specific phenotypes, has led to high levels of homozygosity, and a large undesired drift of alleles causing Mendelian recessive diseases [32, 33]. Accordingly, genetic testing for specific mutations has been developed, primarily for use by breed associations, veterinarians and dog owners (e.g., [34, 35]). For example, GD are currently monitoring multiple mutations using DNA testing and pedigree tracking, in order to reduce their frequency and impact in their population. The UK Kennel Club also undertakes a similar collation of mutation test results, inferring genotypes via pedigree for multiple breeds, and making them available for breeding purposes [36]. However, for many populations, financial or logistic resources to implement and maintain genetic testing using commercial providers are lacking. These tests currently range in cost from around £50 for a single locus, to around £100 for over 200 health-related tests. As an alternative, WGS-based approaches could potentially capture all disease-related variants by default. In contrast, most variants will not be captured by existing canine SNP arrays due to marker patenting unless specifically designed with associated royalty payments for each variant. Thus, for this purpose, WGS could be useful for capturing these previously identified variants. Notably, some disease-causing variants will not be readily captured with the short-read sequencing technology, but a closely linked loci and homozygosity mapping might suffice for practical breeding purposes [37, 38].

However, the majority of diseases and traits of interest are not Mendelian but are “complex”, associated with a large number of genes of small effect and usually strongly influenced by environmental factors. These are more difficult to manage outside of large breeding programmes since a large number of individuals with both phenotypic and genomic information is required for investigation. Considering service dog populations, the majority of withdrawals are due to unsuitable behaviours, followed by health issues. Thus, complex traits of greatest interest to GD and other service dog organizations are those related to behaviour and common health conditions [39]. Many of these traits have been shown to be heritable, making them candidates for selection and for dissection of individual gene effects. For example, moderate to high heritabilities (PB: pedigree-based, G: genomic) have been reported for behaviour traits, including “trainability” (PB: 0.28, [28]), “non-social fear” (PB: 0.25 [28]; PB: 0.36 [40]), and “concentration” (G: 0.63, [41]), as well as health traits, including hip score (PB: 0.28-0.48 [42, 43]) and atopic dermatitis risk (PB: 0.31 [44]). As discussed above, while SNP array data can be effectively used to identify associated genomic regions with genome-wide association studies (GWAS), WGS has been shown to be advantageous for fine-mapping rare variants and for identification of trait-associated genomic regions previously depleted of markers [45, 46, 47]. Additionally, WGS-based GWAS using datasets containing a diverse set of individuals, such as multi- and cross-breed populations, have been shown to identify more variants and QTL than SNP-array-based GWAS performed either within or across breeds (reviewed in [47]). In the case of GD, given their extensive use of crosses, this characteristic of WGS-based GWAS would enhance their efforts to better understand traits, such as complex diseases and behaviour. Furthermore, there is accumulated evidence for the importance of structural variants, such as indels and copy number variants, for complex traits, like reproduction and health traits; there is greater capacity to detect such variants with WGS data than SNP array data (e.g. [48, 49, 47]). Although low-pass sequencing has lower definition than high-coverage WGS, offering less capacity to discover structural variants, it might still preferable over SNP array genotyping for the future dissection of complex traits in the GD population.

### Wider applications

Using simulation to support strategic decision-making is becoming more and more common, (e.g., [50, 51, 52]), and the various points discussed above may be relevant to other populations. In this study, we reported the application and adaptation of the simulation framework to GD and their specific objectives. The pipeline we created in this study could also be applied to other dog populations, particularly for other working dog populations (reviewed in [53]), and even species. A population similar to dogs in terms of structure, organisation and social context is that of horses. For example, breeding of Thoroughbred horses, the most common racing breed, is a multi-billion dollar industry but no global genomics program is in place to enhance performance or control inbreeding [54]. Breeding decisions are usually made based on lineage prestige and individual performances, leading to decreased genetic diversity and higher incidences of Mendelian diseases over the years [55]. Such a population could gain from genomic analysis, in order to better manage inbreeding and to dissect the genetic basis of key traits and diseases. Our simulation protocol could be adapted to evaluate genotyping strategies for Thoroughbred horses. Furthermore, this framework could also be applied to small livestock breeds, where, like GD, the costs of genotyping may be restrictive [56, 57].

The simulation framework developed in this study is highly flexible and each part can be adapted to the specific needs and goals of a given population, making it suitable for many applications. For example, in the case described here, the imputation to WGS was performed with AlphaPeel to take advantage of the pedigree information and pre-existing WGS data, as well as the population structure with large litter and family size in dogs. Other parameters and software for genomic features, initial population, subsequent generations, and imputation can easily be implemented in the framework, which may be more appropriate for other populations. The same framework could also be used to test scenarios based on genomic selection results, by simulating phenotypes and associated genomic predictions, an existing feature of AlphaSimR used in several studies (e.g., [58, 59, 60, 61, 52]).

## Conclusions

This project aimed to develop a simulation framework to determine the best genotyping strategy based on a breeding organisation’s objectives. We based our simulation on the case of Guide Dogs, an organisation managing a small service dog population, which aspires to use genomic information to improve health and welfare of their breeding and working dogs. Simulation was employed to efficiently compare genotyping strategies based on SNP arrays, low-pass sequencing, and high-coverage sequencing, offering insights into imputation accuracy without direct application or testing. From the tested scenarios, low-pass sequencing emerged as the best option by providing high-density genomic information overall and for specific loci of interest, supporting both its breeding goals and genomic research, as well as balancing cost and data processing restrictions. As an alternative, SNP array genotyping would provide adequate information for genomic prediction. However, SNP array scenarios, even supplemented with continued sequencing of breeding stock, had an expected lower imputation accuracy at the whole-genome level and higher cost than low-pass sequencing scenarios for this specific population, resulting in inferior outcomes. The simulation framework developed in this study is highly adaptable and could be modified to address the specific needs of their breeding organisations, thereby offering strategic insights for optimising the collection of genomic information.

## Ethics approval and consent to participate

Not applicable

## Consent for publication

Not applicable

## Availability of data and materials

Code underpinning the created framework (pipeline for simulation, imputation and results processing) is available at https://github.com/HighlanderLab/amartin born2guide seq.The original data (anonymised pedigree) is property of the Guide Dogs for the Blind Association and can be made available upon a reasonable request.

## Author’s contributions

DC, GG, JS, and PW secured funding for the study. TL provided GD data and information. AM and GG designed the simulation pipeline, with consultation from all co-authors. AM performed the simulation, imputation, and results processing. AM drafted the manuscript with feedback from all co-authors. All authors read and approved the final manuscript.

## Author details

^1^The Roslin Institute and The Royal (Dick) School of Veterinary Studies, University of Edinburgh, Easter Bush, EH25 9RG, Roslin, UK. ^2^Hospital for Small Animals, The Royal (Dick) School of Veterinary Studies and the Roslin Institute, University of Edinburgh, Easter Bush, EH25 9RG, Roslin, UK. ^3^The Guide Dogs for the Blind Association, Hillfields, Burghfield Common, Reading, RG7 3YG, Berkshire, UK.

## Additional Files

Additional file 1 — Breed partitioning of the current Guide Dog population.

Dogs included in “Other breeds” can be from one breed (22 other pure breeds) or various crosses.

Additional file 2 — Frequency of sequencing coverage obtained for the current breeding population. Histogram of sequencing coverage for the current GD breeding dogs.

Additional file 3 — Individual imputation accuracy for baseline scenarios.

The whole population is sequenced at 20X coverage and only the current breeding population (Gen 0) previously sequenced has genomic information for None. The brackets represent the 95% interval for the variation observed across the five simulated replicates.

Additional file 4 — Marker imputation accuracy for the region of interest (mutation) for SNP array genotyping strategies.

The brackets represent the 95% interval for the variation observed across the five simulated replicates.

Additional file 5 — Marker imputation accuracy for the region of interest (mutation) for low-pass sequencing strategies.

The brackets represent the 95% interval for the variation observed across the five simulated replicates.

Additional file 6 — Individual imputation accuracy for low-pass sequencing strategies with or without resequencing. The brackets represent the 95% interval for the variation observed across the five simulated replicates.

