## Supplementary figures and images for "Evaluating genotyping strategies for a small managed population with simulation"

### Additional File 1

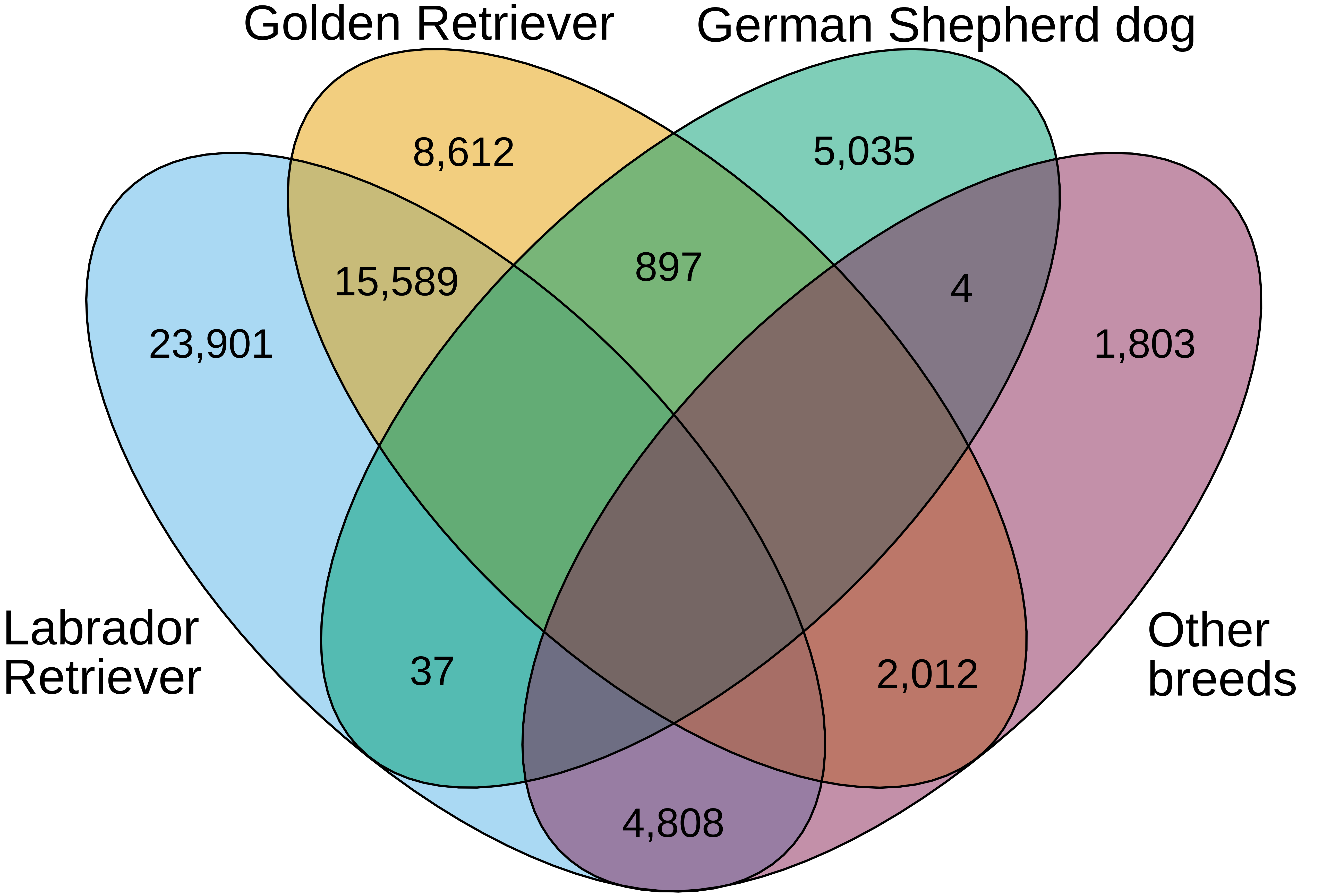

### Additional File 2

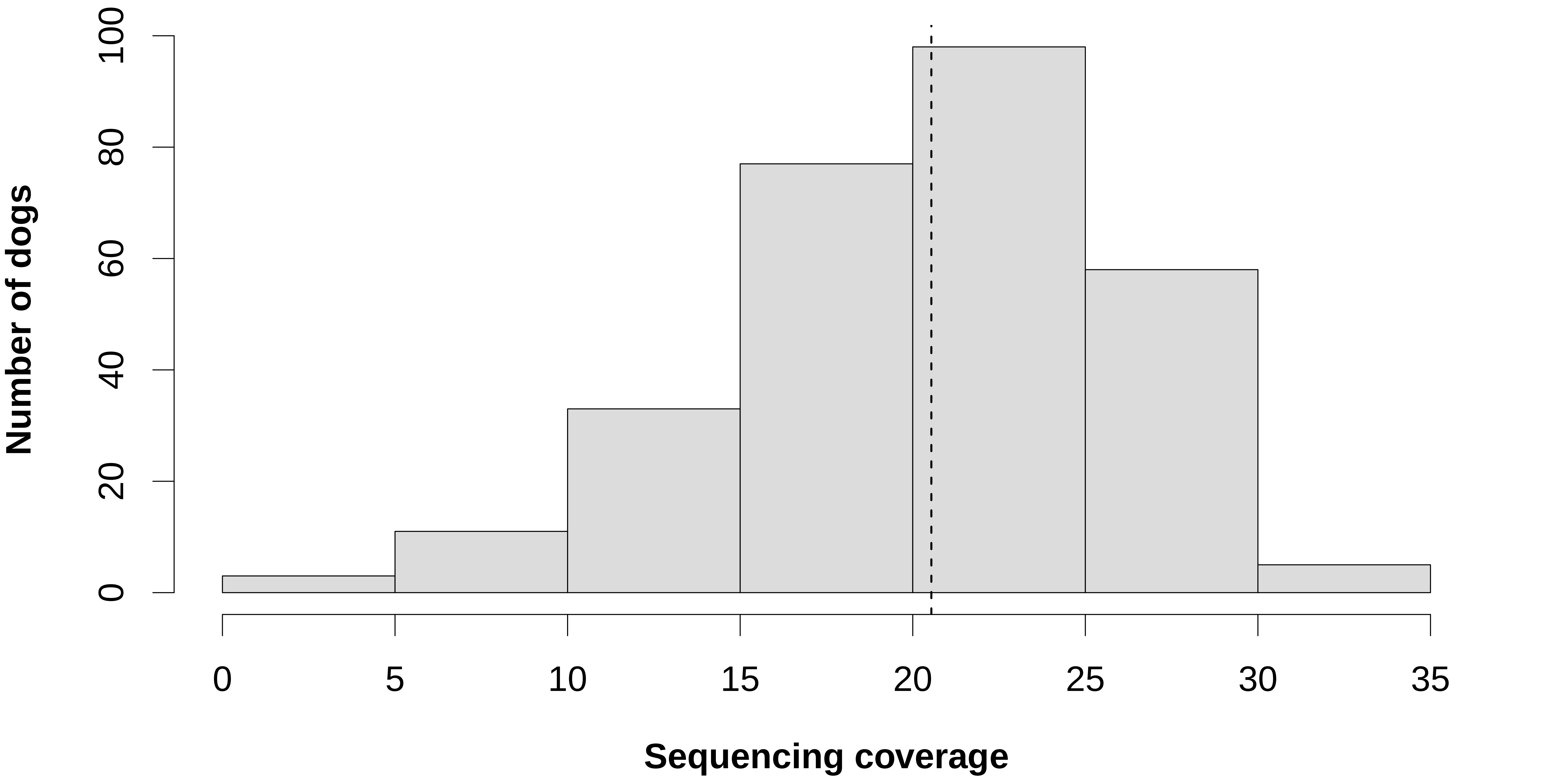

### Additional File 3

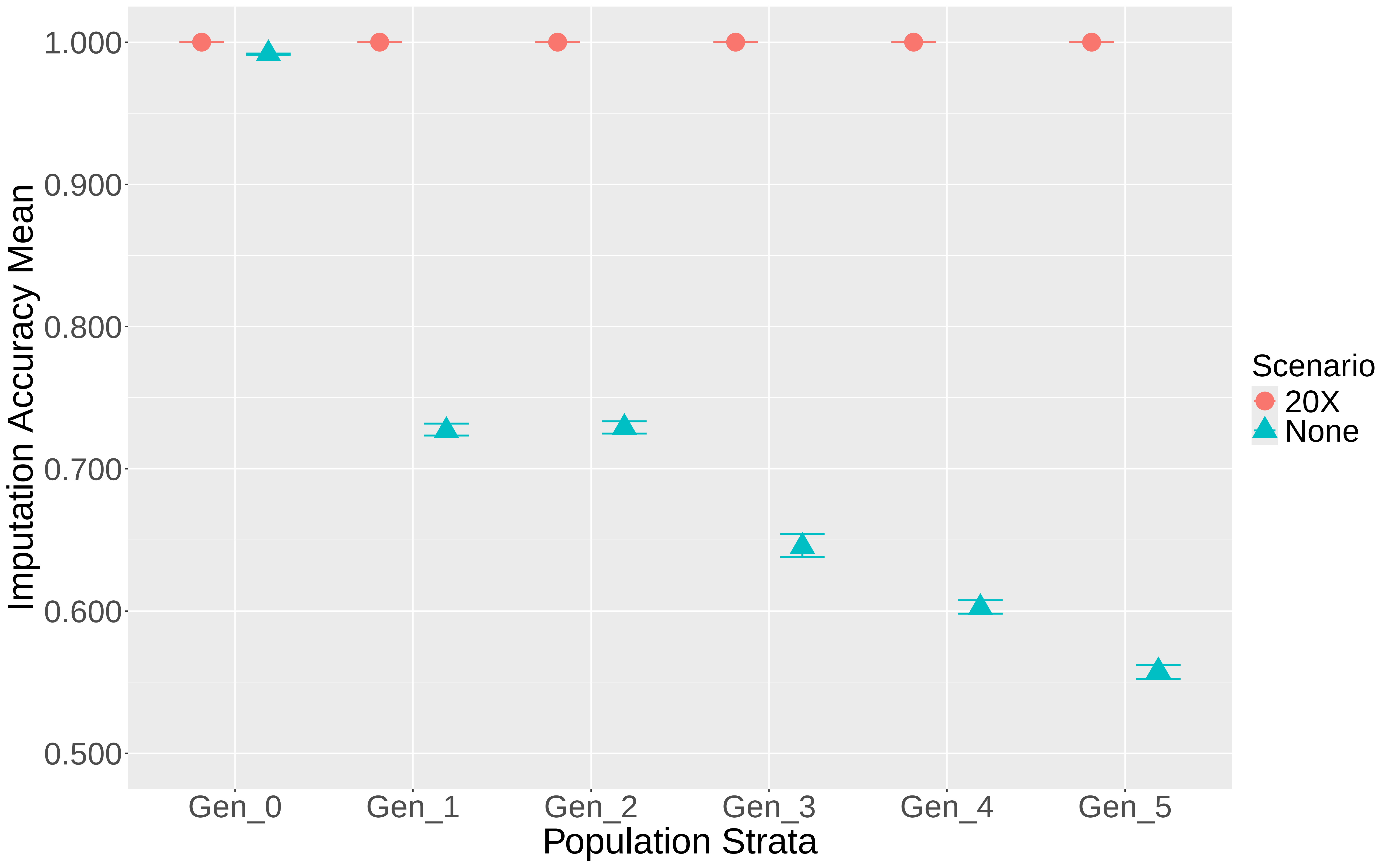

### Additional File 4

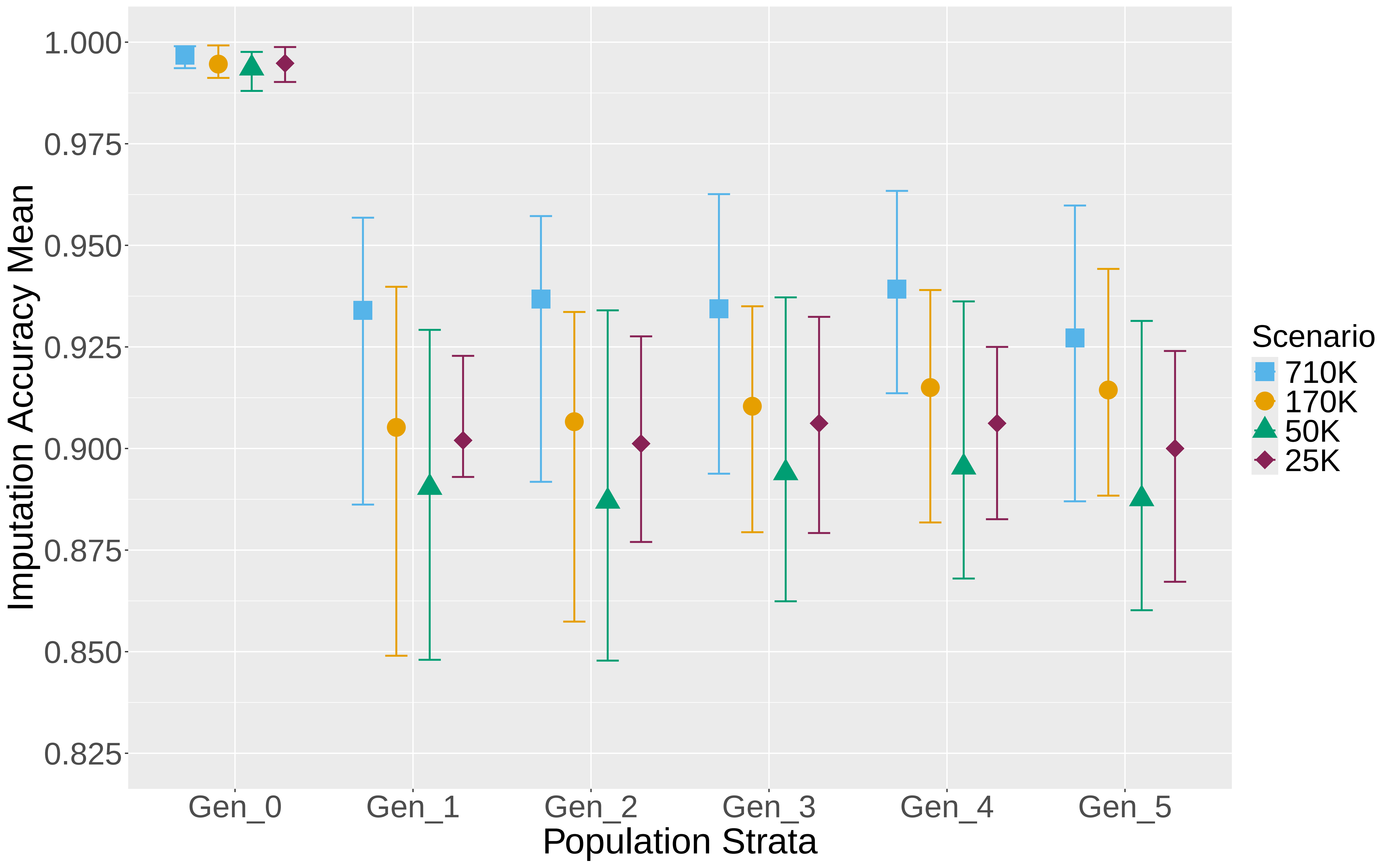

### Additional File 5

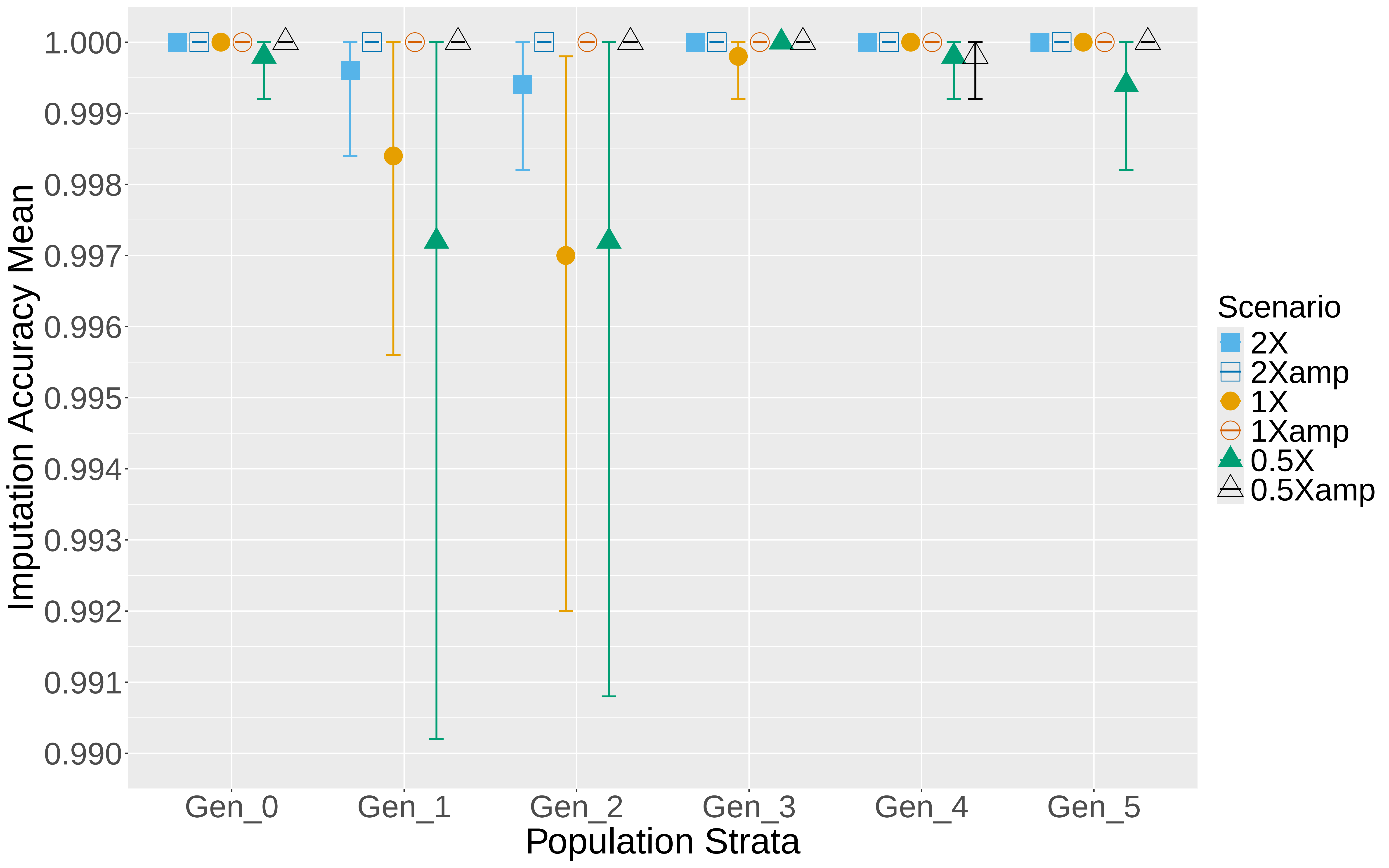

### Additional File 6

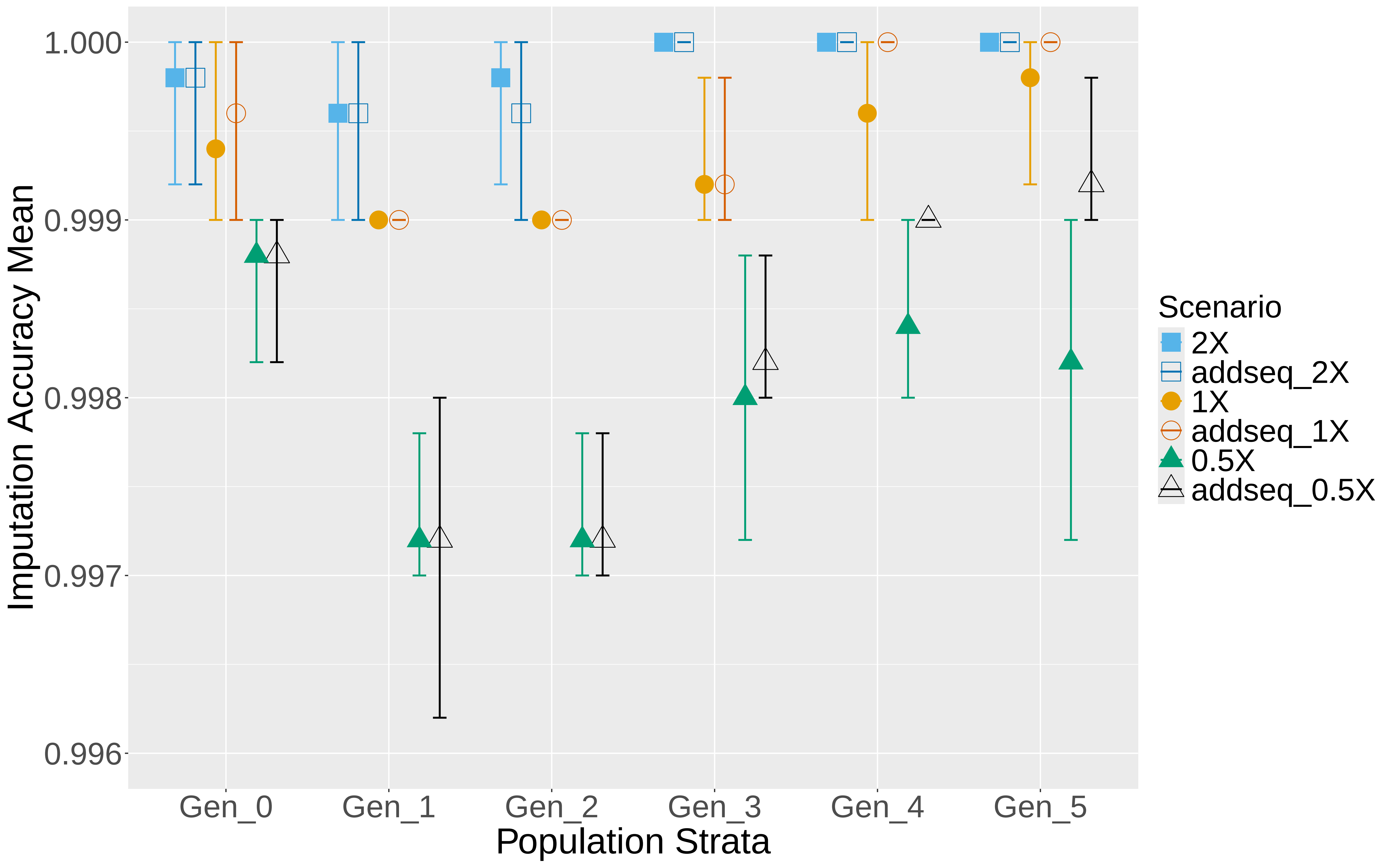
